# Using machine learning to design adeno-associated virus capsids with high likelihood of viral assembly

**DOI:** 10.1101/2021.05.18.444607

**Authors:** Cuong T. To, Christian Wirsching, Andrew D. Marques, Sergei Zolotukhin

## Abstract

We study the application of machine learning in designing adeno-associated virus (AAV) capsid sequences with high likelihood of viral assembly, i.e. capsid viability. Specifically, we design and implement Origami, a model-based optimization algorithm, to identify highly viable capsid sequences within the vast space of 20^33^ possibilities. Our evaluation shows that Origami performs well in terms of optimality and diversity of model-designed sequences. Moreover, these sequences are ranked according to their viability score. This helps designing experiments given budget constraint.

## 1 Introduction

AAVs are the most used viral vectors for gene therapy and considered safe [13]. Nevertheless, it remains a challenge to improve its different properties such as immunogenicity or transduction efficacy. Directed evolution [1] has been used to engineer new recombinant AAVs [5, 8], or more concretely, to design their capsid protein. This is an iterative process which mimics how nature evolves. A library of AAV variants is created by either mutagenesis on one AAV serotype or shuffling of the AAV capsid gene among different serotypes. This library is then subjected to a selection pressure to screen for their function. The top performing variants are selected for the next round of mutagenesis or capsid shuffling. This process is repeated until selected AAV variants satisfy optimization requirements. These variants could then be validated further either in animals or clinical studies.

Unfortunately, the traditional directed evolution has three shortcomings. First, the process does not exploit all the experimental data; the information about the performance of non-selected variants are discarded in each iteration. Second, directed evolution typically starts from a recombinant AAV and a library is built with few mutations per variant. This causes the optimized variants remaining to be similar to the original recombinant AAV. Hence, they are likely to be suboptimal. Finally, because of the nature of experiments, directed evolution is time-consuming and costly. This limits the number of experiment rounds that can be done.

Machine learning (ML)-guided directed evolution [2, 10, 14] could be a solution for the aforementioned problems. We can use the screening dataset to train a predictive model to approximate the mapping from protein sequences to their function, aka fitness landscape [10]. This predictive model can screen virtually variants to save time and cost of experiments, assuming that the approximation is good enough. The virtual screening results are used by model-based design algorithms [3, 9, 12], which explore the sequence space to find optimal sequences. These algorithms are aptly called *explorers*.

In [7], the authors train ML models to predict whether an AAV2 capsid sequence will form a virus particle once produced. The idea is that, with an accurate predictive model, researchers can virtually screen any capsid sequence for its viability without performing actual experiments. This enables them to build better libraries quickly.

While a predictive model helps filtering out non-viable sequences, it does not recommend which sequences to try out to begin with. This is a difficult problem due to the unimaginably large size of the sequence space. In [7], the number of mutational position is 33, which means the corresponding sequence space cardinality is 20^33^! We definitely need smart algorithms to design de novo protein sequences.

In the remainder of this report, Section 2 discusses our approach in detail. Section 3 presents our evaluation. Finally, Section 4 discusses possible next steps.

## 2 Our approach

First, we define some notations and introduce our approach of using model-based optimization [2, 3, 9, 12], where a distribution of protein sequences conditioned on desired phenotypes is computed. Second, we adapt this approach to solve the problem of designing viable AAV2 capsid.

### 2.1 Notation

Let 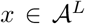 denote an input sequence, where 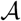 and *L* are the alphabet and the sequence length, respectively. For example, the alphabet of amino acids is {*I,L,...,T,P*}. *h*(*x*) denotes an oracle that predicts the phenotype of a given sequence, *ŷ* = *h*(*x*). Γ ⊂ ℝ denotes the requirement set that characterizes the desired phenotype. For example, in optimizing GFP protein, Γ = *y_fluorescence_* ≥ *y*\*, where *y*\* is a scalar threshold.

### 2.2 AAV2 viral assembly problem

With the defined notation, we view the protein design problem as one of computing a distribution over the protein sequence space conditioned on the desired phenotype, *p*(**x**|Γ). Since we want our AAV2 capsid sequences to be viable, our requirement set takes the form Γ = {*y* =1}. The conditional distribution *p*(**x**|Γ) needs to be computed, and then desired sequences are sampled from it. Let us rewrite *p*(**x**|Γ) as follows.

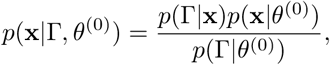

where *p*(Γ = {*y* = 1}|**x**) can be computed by an oracle. *p*(**x**|*θ*^0^) is a generative model that can be learnt from AAV2 capsid sequences. Unfortunately, it is not feasible to compute this condition distribution exactly as the denominator *p*(Γ|*θ*^0^) = ∫*p*(Γ|**x**)*p*(**x**|*θ*^(0)^)*d***x** is intractable due the the high dimensionality of the sequence space.

To overcome the aforementioned intractability, we use the Cross Entropy Method [11], which produces a sequence of generative models, *p*(**x**|*θ*^(*t*)^). Specifically, at each step *t*, it solves the optimization problem:

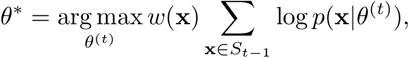

where *S*_*t*−1_ = {**x**|**x** ~ *p*(**x**|*θ*^(*t*−1)^)} and *w*(**x**) weighting function. A popular choice is *w*(**x**) = 1|**x** ∈ *U*_*t*−1_, where *U*_*t*−1_ is a set of sequences whose scores, predicted by the oracle, are above a threshold.

Once we have the final generative model, *p*(**x**|*θ*^(*t*)^), we can sample from it to generate AAV2 capsid sequences with good viability.

## 3 Evaluation

### 3.1 Dataset

The original CapLib8 data set [7] has about 21M data points of the form (*x, y*), where *x* is a sequence of 33 amino acids. These are mutations on the VP1 capsid protein of AAV2. *y* ∈ {1,0} where 1 and 0 stands for viral assembly and non-viral assembly, respectively.

The CapLib8 data set has a class imbalance between non-viable and viable sequences; there are more viable sequences than non-viable ones. As we will see, this affects the oracle performance with respect to the non-viable capsid sequences.

### 3.2 Oracle

We use an LSTM [4] for our predictive model. We sample from the CapLib8 dataset randomly to create datasets of different sizes. The same LSTM architecture is trained on them to evaluate how the performance varies with the size of training data set.

We notice that the recall score for non-viable capsid sequences improves with more data (Figure 1b). This is important because for Origami to work, we need to prevent training our generative model with false positive sequences, i.e., sequences that are predicted to be viable but are in fact non-viable. The effect of the data set size is not significant for the oracle performance with respect to viable sequences (Figure 1a).

**Figure 1:**
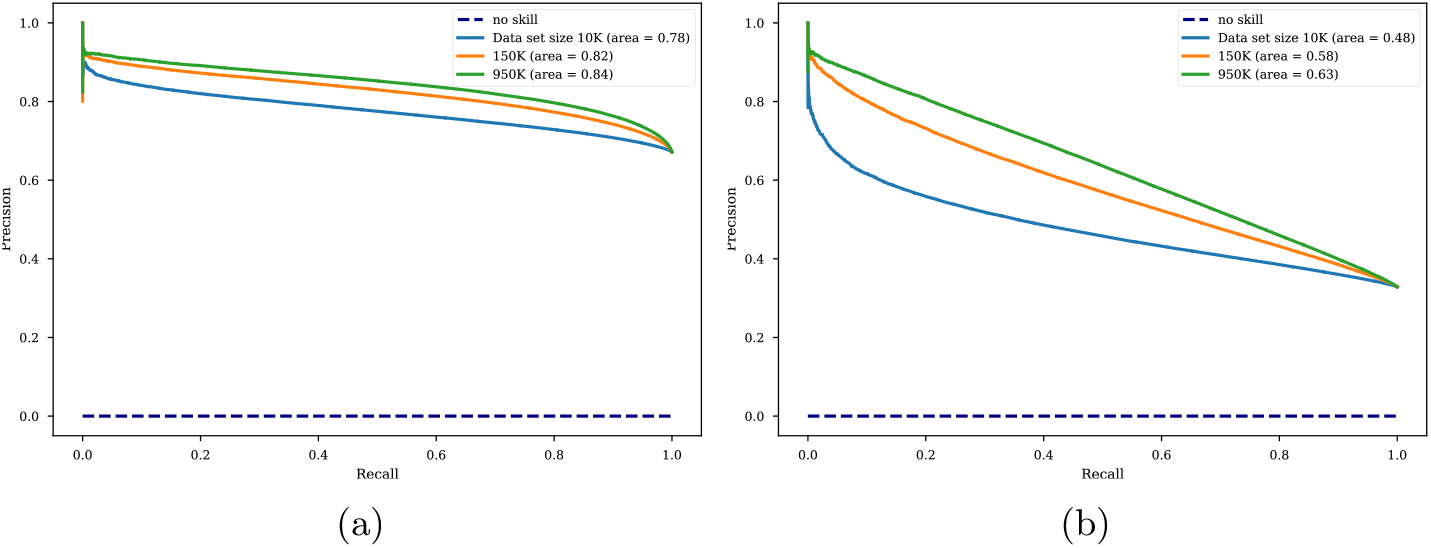
The quality of oracles with different training dataset sizes.(a) Precision recall curve with respect to viral assembly. (b) Precision recall curve with respect to non-viral assembly.

Finally, we create a balanced data set of total 700K data points with equal split between viable and non-viable AAV2 capsid sequences. We use this data set to train an oracle that will be used by our Origami algorithm to screen for capsid viability.

### 3.3 Viable sequences

We train a variational auto encoder (VAE) [6] as our generative model with tuned hyperparameters. To validate Origami, we perform viral assembly predictions on a test data set of 150K data points. Then we take those sequences in the bottom 20th percentile of prediction scores. These sequences are used as the starting sequences for Origami. We run Origami for 20 iterations. At the end, the model-designed sequences have an average assembly likelihood of 0.75, which is 4.7 times better than that of the starting ones (Figure 2b).

**Figure 2:**
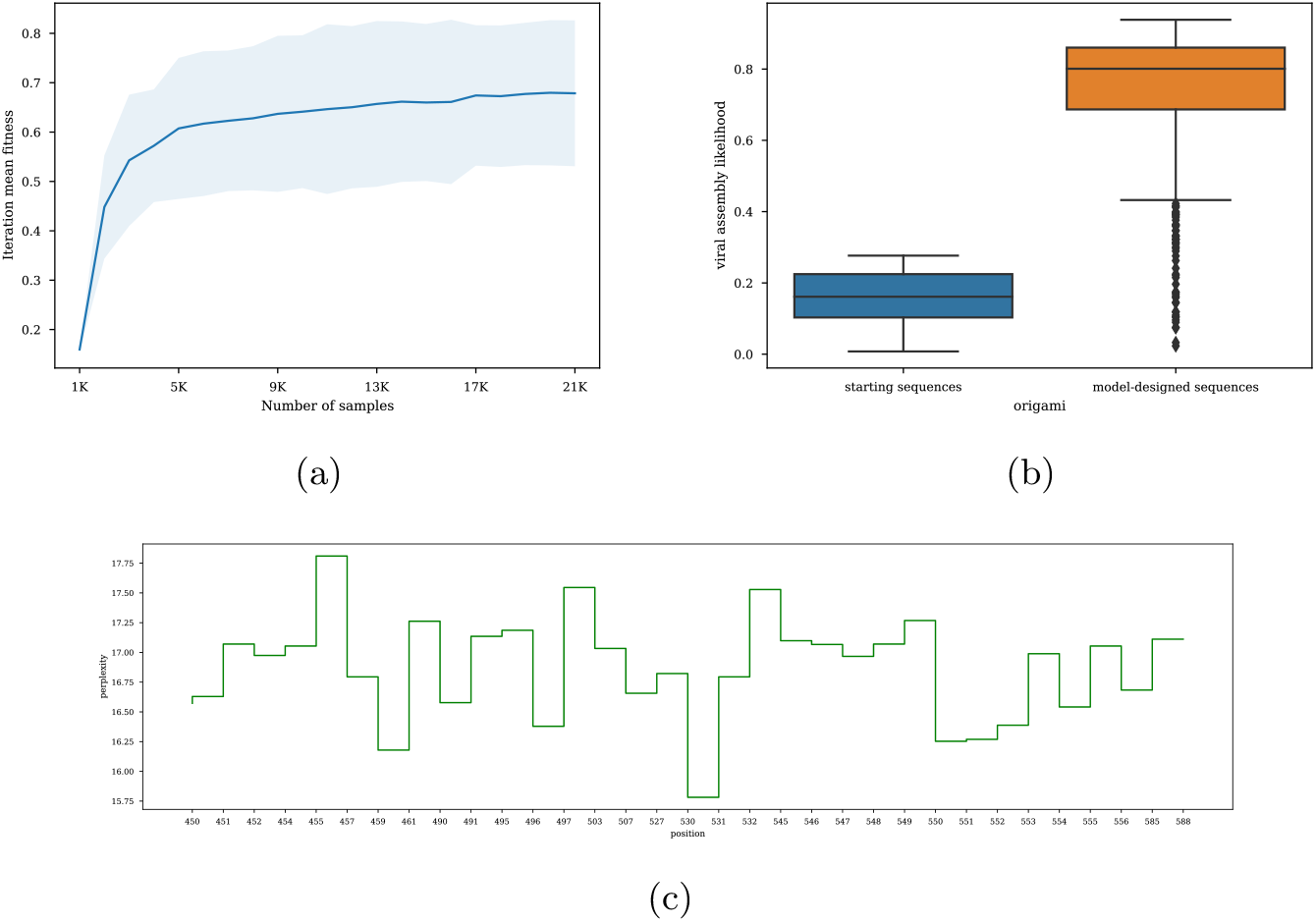
(a) Model-designed sequence mean fitness improves with each iteration. Note that the x-axis shows the cumulative number of samples over iterations. The shade area shows the 95% confidence interval. (b) Starting sequences vs. model-designed ones in terms of viability score, i.e. the probability of viral assembly provided by the oracle. (c) The perplexity of distribution of amino acids at each of the 33 mutational positions. The higher the perplexity, the higher the diversity.

Figure 2c the perplexity at each position of the mutated AAV2 capsid sequence. The perplexity value can be viewed as “branching factor”. In other words, it measures the diversity in terms of amino acids found at each position. We can see that the model-designed sequences are quite diversified.

We also evaluate our optimization procedure using the metric *iteration mean fitness*, which is the average fitness score of the current batch, *S*_*t*−1_ = {**x**|**x** ~ *p*(**x**|*θ*^*t*−1^)}. Figure 2a shows that Origami can generate sequences with increasing viral assembly likelihood over iterations.

## 4 Future work

We must admit that the CapLib8 dataset is not an ideal data set to evaluate Origami. First, as per private communication with authors of [7], to build the CapLib8 dataset, the authors have to sequence the initial library creating a sequence set called 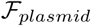, and then after the viral production, sequence it again to create another sequence set called 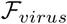. All the sequences in 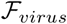 are considered viable whereas all found in 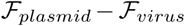 are considered non-viable. Unfortunately 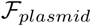 was not fully sequenced, hence CapLib8 does not have a lot of mutants that did not assemble in virus production. This creates a class imbalance that affects the performance of oracles as discussed in Section 3.2.

Second, Origami was designed to optimize under the assumption that sequences with good phenotypes are rare events. But since the “good” event of viral assembly is abundant in the CapLib8 dataset, it is therefore not an ideal one to benchmark Origami.

Finally, CapLib8 provides only one single screening dimension, namely viral assembly. Therefore, we could not evaluate Origami in a multi-objective optimization scenario. The next step could be using a multi-organ bio-distribution data set to evaluate Origami further.

